# No evidence for kin recognition in a passerine bird

**DOI:** 10.1101/560870

**Authors:** Martina Lattore, Shinichi Nakagawa, Terry Burke, Mireia Plaza, Julia Schroeder

**Affiliations:** Department of Life Science, Imperial College London, Silwood Park, Ascot, UK; Evolution & Ecology Research Centre and School of Biological, Earth and Environmental Sciences, University of New South Wales, Sydney, Australia; Department of Animal and Plant Sciences, University of Sheffield, Sheffield, UK; Department of Evolutionary Ecology, National Museum of Natural Sciencie-CSIC, Madrid, Spain

## Abstract

Theory predicts that individuals behave altruistically towards their relatives. Hence, some form of kin recognition is useful for individuals to optimize their behaviour. In species displaying bi-parental care and subject to extra-pair matings, kin recognition theoretically allows cuckolded fathers to reduce their parental investment, and thus optimize their fitness, but whether this is possible remains unclear in birds. This study investigates the ability of male sparrows to recognize their own chicks, using a large cross-foster experiment, parental care as an indicator and House Sparrows (*Passer domesticus*) as a model organism. We cross-fostered chicks after hatching, and then expected that fathers would show a decrease in their parental efforts when tending to a clutch of unrelated offspring. However, there was no significant effect of relatedness on provisioning rates. This suggests that sparrows may not be capable of kin recognition, or at least do not display kin discrimination despite its apparent evolutionary advantage.

## INTRODUCTION

Kin recognition is “the ability to recognize one’s genetic relations” [1] and evidence for it has been found to be widespread across many taxa [2,3,4,5]. Indeed, individuals can behave altruistically [6] and more so the more two individuals are related [7]. Hence relatedness is a determining factor in understanding how altruism works in nature, and therefore a driver of behavioral evolution [8].

Relatedness is the proportion of shared genes between conspecifics. Building upon Fisher’s [9] work on altruistic behavior, Hamilton [7] postulated that an altruistic behavior will be expressed when the costs of the behavior are lower than the fitness benefits to the individual benefitting from the altruistic behavior, modulated by the degree of relatedness. According to Hamilton’s Rule, for a behavior to be considered altruistic the following equation must be achieved:

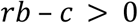

where *r* is the genetic relatedness between individuals, b is the fitness benefit to the receiver, c is the fitness cost to the altruist. Hence, *rb* corresponds to the indirect fitness effects of a trait, whereas -c is the direct effect [10]. For instance, in a parents-offspring relationship, the parent incurs costs, by spending energy finding food, not eating it themselves but feeding it to their young. Yet, they are genetically related to their offspring, and thus the fitness benefit that the offspring received by being fed translates into fitness for the parents – but only as long as they are related to each other. Hence, mechanisms allowing individuals to recognize relatives may be useful, in species where kin-selection is favorable [7].

There is an ongoing debate on the details of the definition of kin recognition, considering different aspects of it, including genetic and environment-based cues and mechanisms. For the purposes of this work, the most relevant definition is the operational one suggested by [11], with kin recognition being defined as the “differential treatment of conspecifics differing in genetic relatedness” (also referred to as “kin discrimination” [1]).

There are numerous mechanisms for individuals to recognize their genetic relatives. Contextual cues include spatial and temporal factors, and this mechanism is quite common among animal species [8]. Birds, in particular, often seem to recognize any chick in their nest as their own, however they can also take into account their access to a mate and the probability of their paternity of the brood to judge whether the young is theirs [12]. Instead, kin recognition via phenotypic cues takes into consideration a variety of phenotypic traits of the individuals. The “prior association hypothesis” is based on direct familiarity, where individuals recognize their own kin by first becoming familiar with their phenotypes in a shared, early environment and then recognizing them in the future [8, 11]. The “phenotype matching hypothesis” suggests that “individuals who resemble their own kin are treated as related” [8].

However, the ability to recognize their own kin can enable individuals to adjust their behavior according to the degree of relatedness to others [1, 11]. In species that display biparental care such as many social birds, it is thought the father’s effort might be affected by his relatedness to the nestlings if he is capable of recognizing them [13]. Caring for unrelated offspring has no positive effect on an individual’s fitness as they would be spending unnecessary effort and resources to raise offspring that does not share any of their genetic material [14]. Caring for unrelated young is therefore considered an unnecessary cost that is best avoided or reduced [15]. As females are typically fairly sure of their genetic relatedness to their young, fathers experience a greater uncertainty of paternity, and thus, one would expect it to be beneficial if fathers could distinguish between kin and non-kin. It is therefore expected that males reduce their parental care to young they are not related to.

Some experimental and observational research on kin recognition has produced contrasting results [16, 17, 18, 19], due to different risks of extra-pair offspring and other ecological factors in the studies. These suggest that overall, birds do not differentiate between relatives and non-relatives. For example, Boncoraglio and Saino [18] showed that Barn Swallows (*Hirundo rustica*) fathers do not underfeed unrelated chicks, even though chicks themselves beg differently when in mixed broods. In contrast, Zebra Finches (*Taeniopygia guttata*) appear to behave differently towards related and unrelated offspring [19].

Because of this limited knowledge on the mechanism of kin recognition in birds, we here test the hypothesis that some birds are capable of kin recognition in a large, long-term dataset with information on genetic relatedness in House Sparrows (*Passer domesticus*).

This social bird species has a relatively high extra-pair paternity rate [20, 21], which makes house sparrows an ideal model to investigate kin recognition in. Female infidelity has a negative effect on males’ fitness, as males receive no fitness benefit from caring for chicks that do not share their genes [15]. Male individuals would therefore be expected to provide less parental care to nestlings born from extra-pair matings by their female. This however requires the males to be able to recognize his kin recognition to adjust their level of care.

## METHODS

### Study population

The house sparrow is a social bird with bi-parental care. Adults form social pairs that last for one or multiple breeding seasons. However, while socially monogamous, there is some genetic infidelity present in this species [21]. Furthermore, males seem to employ a bet-hedging strategy where they adjust their parental care according to the female they are paired with, and while there is a genetic basis, most of the within-male variation is due to phenotypic plasticity or unexplained [22, 23]. Thus it is a suitable species to study kin recognition between parents and offspring.

This research was carried out on a population of house sparrows that has been monitored since 2000 on Lundy Island, 19km off the coast of Devon in the Bristol Channel (51°10′N, 4°40′W). where Levels of migration reach a maximum of three birds every four years [20], making it a nearly closed population suitable for longitudinal studies. Life histories of individual birds and a full pedigree of the population are available, therefore the identities of parents for each brood are known [24]. Birds are individually marked with a unique combination of three colored rings and a numbered metal ring. In addition, each sparrow was provided with a subcutaneous passive integrated transponder which is read by radio frequency identification (RFID) antennas attached to each nest-box on the island [25]. A total of 95% of the breeding occurs within these nest-boxes. All of them were monitored both by direct observations and using cameras as follows [26].

### Data collection

We used data collected from chicks born between 2004 and 2015, as part of the Lundy Island Sparrow Project. The chicks were cross-fostered on the day after hatching and remained in their rearing brood until fledging. Each brood underwent one of three treatments, where the brood was either not cross-fostered, partially cross-fostered, or fully cross-fostered [27]. However, we had information about genetic paternity in all broods, and we used that to consider the status of the father in relation to the brood it attended. Using all this information, we created a variable with three treatment levels with respect to the genetic relatedness to the social father who attended the brood: fully related (−1), partially related (0) and unrelated (1).

All broods were monitored with video-cameras placed between 2 and 5 meters away from their entrance, with a field of view of 30cm around the nest-box [26]. Videos were recorded in the morning between day 1 and day 15 after hatching. This study uses the time from when a bird is first seen on video to the end of the recording as “effective video time”, to account for the fact that individuals needed time to adjust to presence of a camera in front of their nests [26]. Effective video times range between 10 and 120.26 minutes, with a mean of 88.34 minutes. By watching the videos we extracted the number of parents visits to the nest and we calculated the provisioning rate for each parent (number of visits per hour). Only the provisioning rate of the social father was used in this research.

### Statistical analysis

We analyzed the data with R version 3.5.2 Eggshell Igloo [28]. We ran a linear mixed effects model to test the hypothesis that male birds reduce their provisioning rate when tending to broods that hold chicks they are not related to, with the number of provisioning visits per hour by the attending male as the response variable, and parental care as explanatory variable (see below). The standardised age of chicks at the time of the video and the number of hatchlings in a nest were used as additional fixed covariates, as we know they are related to parental care [29]. We also included the brood ID as additional random effect in order to account for the repeated measurements in broods over several years. We used the within-subject centering method [30] to distinguish between a bias of cross-fostered nests towards nests with attending males with higher or lower parental care, and *vice versa*. Such a bias could come across if cross-fostering cannot always happen completely randomly [27], due to cross-fostering always requiring at least two broods of the same age. This means that by definition, broods that are less synchronized with others, such as the very early or very late broods, are unlikely to be cross-fostered. The parents tending to these nests are systematically different from those that breed during time periods when most of the population breeds, as early birds may be of higher quality, and late birds may be of lower quality, on average. To distinguish between a male displaying different parental care towards broods with young that are related to him *versus* a brood with young unrelated to him, and differences in parental care between different male individuals that tend to nests that are cross-fostered and not, we created two new fixed covariates from the variable treatment (−1 = genetic sire to all offspring in the nest, 0 = attending male is related to part of the offspring in the nest, either through partial cross-fostering or through extra-pair paternity, 1 = attending male is unrelated to the offspring in the nest through full cross-fostering). To estimate and capture the between-male variation, we calculated the average value for each male of the treatment. Thus, if a male twice tended to a fully cross-fostered brood (both with a value of 1), and once for a brood containing only his genetic offspring (with a value of −1), this between-male treatment effect (T_b_ in the following) would be 0.33 for each of these observations. To estimate the within-male variation, we subtracted the mean treatment of each male from the treatment value of each observation of every respective individual male. Thus, the male from above would get a value of 0.67 for the first two broods, and −1.33 for the second one for the within-male treatment effect (T_w_ in the following). We used the attending male’s ID as a random effect on the intercept.

The model was run in MCMCglmm (Hadfield 2010). We present parameter estimates with their respective 95% credible intervals (95CI). We considered fixed effects as statistically significant if their 95CIs did not span 0. We also present the pMCMC, that is approximately twice the MCMC estimate of the probability that the 95CI does not span 0, a value analogue to traditional *P* values.

## RESULTS

### Descriptives

We used 2388 video observations collected between 2004 and 2015. We extracted parental care rate for the male from each video, which were from 299 individual male sparrows. Only 19 males were observed only once, all other males were observed repeatedly (Table 1). The median number of times a male was observed was 5, the mean was 7.98 times. Our data was also structured by brood. Of the 1048 broods, 164 broods were only observed once, with an overall median of each brood being observed two times (Table 1).

**Table 1:**
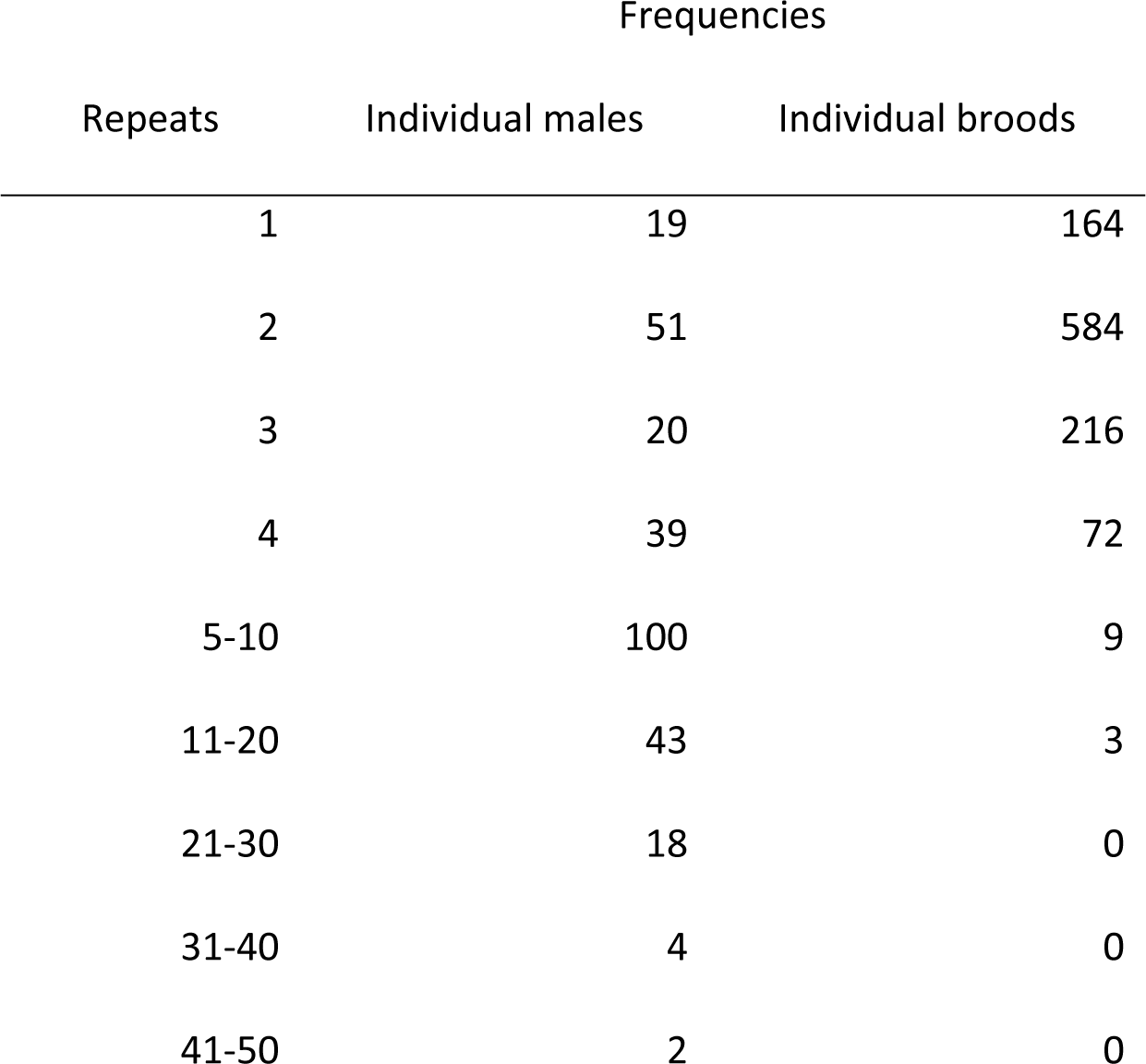
The frequencies of individual repeated observations of parental care of Lundy sparrow males tending to their broods between 2004 and 2015.

| Repeats | Frequencies |  |
| --- | --- | --- |
|  | Individual males | Individual broods |
| 1 | 19 | 164 |
| 2 | 51 | 584 |
| 3 | 20 | 216 |
| 4 | 39 | 72 |
| 5-10 | 100 | 9 |
| 11-20 | 43 | 3 |
| 21-30 | 18 | 0 |
| 31-40 | 4 | 0 |
| 41-50 | 2 | 0 |

Of all 1048 broods, 394 were from non-cross-fostered broods, 196 from fully cross-fostered broods and 458 from partially cross-fostered broods. After accounting for the genetic relatedness of the offspring present in the nest in relation to the attending male, we had 227 broods attended by a male who was related to all offspring in the brood, 591 broods in which some of the chicks were related to the attending male, and 230 broods where the male was completely unrelated to the chicks he was caring for.

Males tended to attend to broods completely related to them with 8.9±0.3 (mean ± SE) visits per hour on average, to partially related broods with 8.8±0.3 visits per hour, and to completely unrelated broods with 8.6±0.4 visits per hour (Fig. 1).

**Figure 1:**
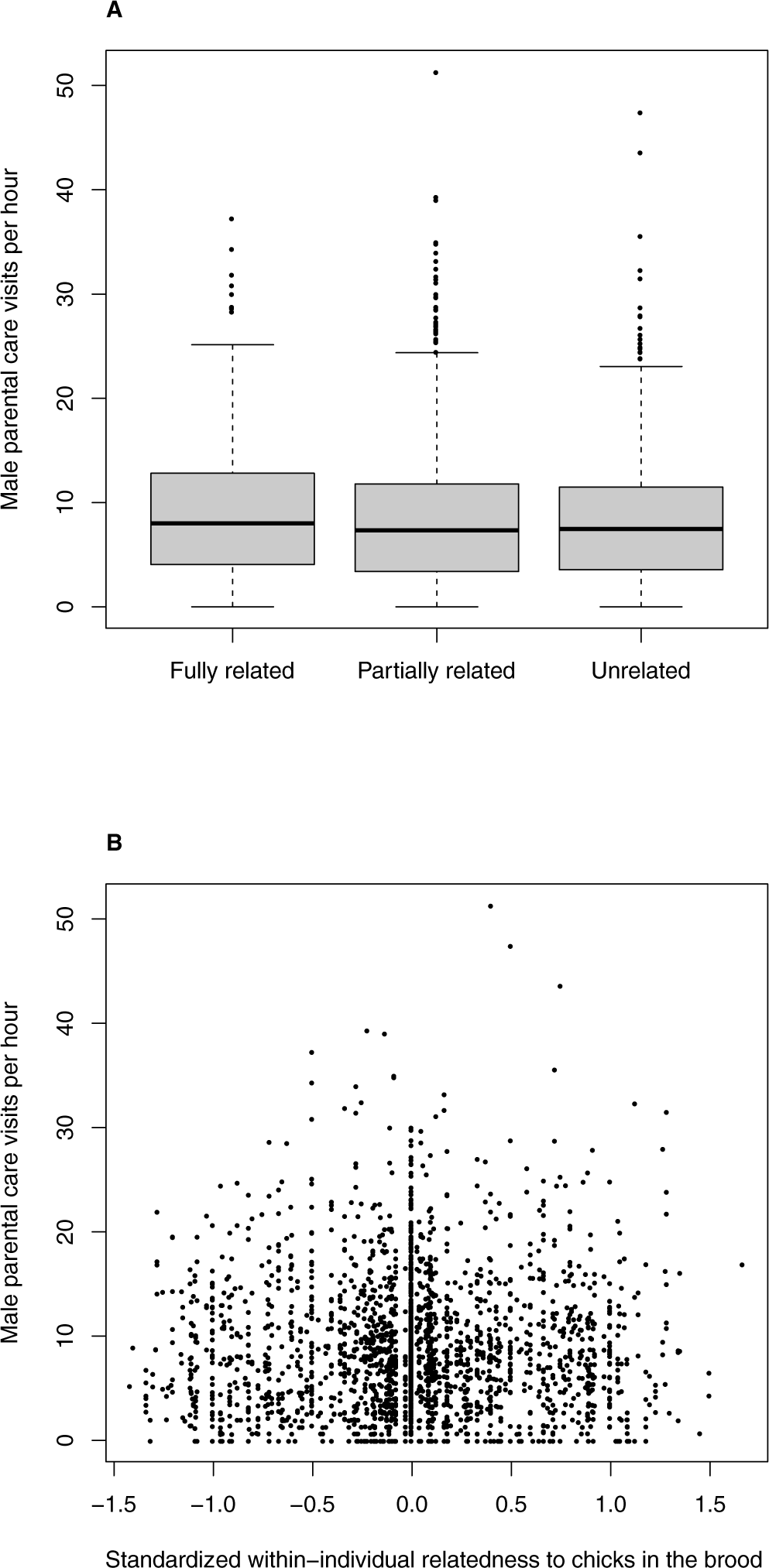
A Male parental care per hour in relation to his relatedness to the brood he cares for. B Male parental care in relation to the difference between an observation and the average relatedness of the respective male to the chicks he cares for. This standardized variable represents variation within individuals. Larger values represent a lower relatedness between the male and his brood. The hinges represent the first and third quartiles, the thick line is the mean, and the dots represent single observations outside of the hinges.

### Do males care less for unrelated offspring?

Males did not change their parental care when caring for a brood containing chicks that were not genetically related to them (Table 2).

**Table 2:** Results from a linear mixed model. Chick age and the number of hatchlings were standardized to a mean of 0 and variance of 1. The treatment refers to the degree of which the chicks were related to the attending male (low = high relatedness, high = low relatedness). N = 2388 observations of parental care between 2004 and 2015, on 1048 broods and 299 individual males.

| *Parameter* | *Estimate* | *Precision* | *Probability* |
| --- | --- | --- | --- |
| *Fixed effects* | *B* | *95CI* | *pMCMC* |
| Intercept | 8.80 | 8.37–9.25 | <0.001 |
| Within-individual treatment | 0.19 | -0.37–0.75 | 0.50 |
| Between-individual treatment | -0.86 | -1.18–0.16 | 0.10 |
| Chick Age | 0.32 | 0.11–0.54 | 0.002 |
| Number of hatchlings | 0.38 | 0.09–0.68 | 0.01 |
| *Random Effects* | *Variance* | *95CI* |  |
| Brood reference | 12.74 | 9.97–15.47 |  |
| Male ID | 6.07 | 3.65–8.46 |  |
| Residual | 24.85 | 22.92–26.76 |  |

## DISCUSSION

We found no support for the idea that male house sparrows recognize their kin and react to this by a reduction of paternal care. Little research has previously been done on kin recognition and paternal care in house sparrows [29].

House sparrows have been shown in the past to be capable of paternal care adjustment, as males with cheating partners show reduced parental care investment [29]. Here, we specifically investigated whether this adjustment was due to the amount of unrelated offspring being present at the nest. If an adult male is able to recognize its own chicks we would expect to see different behavior at nests where we cross-fostered the chicks and where we did not.

This study shows that males do not decrease their provisioning rate towards cross-fostered (or otherwise unrelated) chicks, despite the apparent evolutionary advantage of paternal care adjustment to unrelated offspring. This experimental result, with a high statistical power, does not support the idea that male sparrows are able of kin recognition and it is thus consistent with previous empirical findings. Previous studies have not specifically manipulated conditions for paternal care adjustment in relation to kin recognition before [17, 31, 32]. While our study did not conclusively rule out kin recognition (males could recognize their kin but not react to this information), it is, to our best knowledge, the largest experimental dataset in the wild that is currently available.

It has previously been suggested that trade-offs might limit the advantage of kin discrimination. As one can imagine that kin recognition might not be perfect, the balance between the costs of wrongly rejecting offspring and the costs of wrongful acceptance will play an important role [8, 13, 18, 29]. Furthermore, only part of the chicks in a brood may be unrelated, while another part is related, and a reduction in parental care may be difficult to target at certain chicks only. Moreover, if females seek extra-pair matings to improve their reproductive output, especially by increasing the quality of the genes of their offspring, they would have interest in their extra-pair offspring growing [13]. Therefore, if the males reduced feeding their parental investment, females might increase their own investment. The interplay between male and female parental care is complex, as it is guided by sexual conflict, cooperation, compatibility and several phenotypic, and environmental, indirect social effects [22].

Another explanation for our findings is that kin recognition might derive from learned cues, which require longer to develop than the two days parents spent with their own chicks in this experiment before cross-fostering [31]. For example, Razorbills (*Alca torda*) accept unrelated chicks in their nest and raise them without distinctions within the first 15 days of their lives, whereas if foreign chicks are introduced after 16 days they get rejected [33]. The same phenomenon was observed in European Starlings (*Sturnus vulgaris*), with foreign chicks over 16 days old not being accepted into unrelated broods [34]. The sparrows on Lundy island were a maximum of two days old when cross-fostered (because sometimes not all eggs hatch on the same day), which might have affected the fathers’ ability to recognise them, not being given time to learn their phenotypic cues. If this was the case, it would suggest kin recognition occurs by prior association [8, 11] Future studies using cross-fostering of chicks at a later age could be useful to investigate this.

In summary, this long-term experiment, providing a large amount of data, did not provide support for kin discrimination in the house sparrow. More work is needed to conclusively test whether birds able to recognize their offspring.

